# Genome sequence and annotation of *Periconia digitata*, a promising biocontrol agent of phytopathogenic oomycetes

**DOI:** 10.1101/2023.03.17.533154

**Authors:** Elena Bovio, Corinne Rancurel, Aurélie Seassau, Marc Magliano, Marie Gislard, Anaïs Loisier, Claire Kuchly, Michel Ponchet, Etienne G.J. Danchin, Cyril Van Ghelder

**Affiliations:** Institut Sophia Agrobiotech, INRAE 1355, CNRS and Université Côte d’Azur, 400, Route des Chappes, BP 167, 06903 Sophia Antipolis Cedex, France; GeT-PlaGe (genomic platform), Campus INRAE, 24 chemin de borde rouge, Auzeville CS 52627, 31326 CASTANET-TOLOSAN Cedex, France

**Author notes:** The two first authors contributed equally to this work (Elena Bovio, Corinne Rancurel). Corresponding author: Elena Bovio, Corinne Rancurel.

## Abstract

The *Periconia* fungal genus belongs to the phylum Ascomycota, order Pleosporales, family Periconiaceae. *Periconia* are found in many habitats but little is known about their ecology. Several species from this genus produce bioactive molecules. *Periconia digitata* extracts were shown to be deadly active against the pine wilt nematode. Furthermore, *P. digitata* was shown to inhibit plant pathogenic oomycete *Phytophthora parasitica*. Because *P. digitata* has great potential as a biocontrol agent and high quality genomic resources are still lacking in the Periconiaceae family, we generated long-read genomic data for *P. digitata*. Using the PacBio Hifi sequencing technology, we obtained a highly-contiguous genome assembled in 13 chromosomes and totalling ca. 39 Mb. In addition, we produced a reference transcriptome, based on 12 different culture conditions, and proteomic data to support the genome annotation. Besides representing a new reference genome within the Periconiaceae, this work will contribute to our better understanding of the Eukaryotic tree of life and opens new possibilities in terms of biotechnological applications.

## Introduction

In 1934, the family Periconiaceae was established with *Periconia* as type genus (Hyde et al., 2018). Subsequently, species belonging to the genus *Periconia* have been assigned to the family Massarinaceae, but a recent systematic revision brought all the *Periconia* spp. in the family Periconiaceae (Tanaka et al., 2015). The *Periconia* genus was established in 1791 by Tode Ex Fries (Hyde et al., 2018), while the first record is dated back to a fossil preserved in the Baltic amber from the upper Eocene epoch (34-38 Mya) (Tischer et al., 2019). The genus comprises 211 species epithets in Index Fungorum (2022) with 25 of them having a new current name belonging to another genus. To date, only 27 species have been confirmed by molecular data (Yang et al., 2022). The habitat of *Periconia* is various, the reports start from the sea (Bovio et al., 2018; Cantrell et al., 2007) to the Himalayas (Prasher & Verma, 2012). To the best of our knowledge, most species present a saprobe behaviour (Crous et al., 2018; Hyde et al., 2018; Markovskaja & Kačergius, 2014; Yang et al., 2022), while some are described as endophytes and are included in the dark septate endophytes - DSE group (J. Y. Li et al., 1998; Mandyam et al., 2012; Verma et al., 2011). Some endophytic *Periconia* spp. were found to be facultative parasites on the invasive weed *Parthenium hysterophorus* (Asteraceae) (Romero et al., 2001). *P. circinata* was described as the causal agent of root rot on *Sorghum* spp. (“milo disease”) (Odvody et al., 1977). *Periconia* spp. were also found associated with human keratomycosis (Gunasekaran et al., 2021). Although the genus *Periconia* is present in many habitats, little is known about its ecology. Many studies only focused on the ability of given species (not always identified at species level) to produce molecules of biotechnological interest. Since 1969, the genus has yielded 104 compounds belonging to terpenoids, polyketides, aromatic ketone and phenolics, some exhibiting interesting biological activities including antimicrobial (bacteria, fungi), antiviral, anti-inflammatory and cytotoxic activity (Azhari & Supratman, 2021). The anti-oomycetes activity of a *Periconia* (strain Y3) has been explored only once (Galiana, Marais, et al., 2011; Galiana, Ponchet, et al., 2011). The strain Y3 was first misidentified as *Phoma* sp. CNCM I-4278 according to its 18S ribosomal sequence (accession number HM161743). However, taking advantage of the recent systematic revision of the genus *Periconia* and the increasing set of sequences now available (Hyde et al., 2018; Tanaka et al., 2015), the strain CNCM I-4278 was correctly identified as *Periconia digitata* and confirmed by ITS and 28S sequences. *P. digitata* CNCM I-4278 was able to inhibit the growth and cyst germination of the plant pathogenic oomycete *Phytophthora parasitica* both in vitro and in planta, without phytotoxicity (Galiana, Marais, et al., 2011; Galiana, Ponchet, et al., 2011). In addition to anti-oomycete activity, the water filtrate and/or the crude extract of *P. digitata* CNCM I-4278 also inhibited the growth of several fungi that are well known plant pathogens (Galiana et al., 2011; unpublished data). In another screening of fungal culture filtrates isolated from freshwater submerged wood, a *P. digitata* strain was highly active on the fungivorous and phytophagous nematode *Bursaphelenchus xylophilus* responsible for dramatic losses in pine forests. Indeed, 70% to 80% of nematodes were killed 48 h after the treatment with *P. digitata* extracts (Zhu et al., 2008).

Owing to its high potential as a biocontrol agent for plant protection against several classes of problematic plant pathogens, we present the chromosome-scale genome assembly and annotation of *P. digitata* CNCM I-4278. To date, only 2 other genomes were available in the genus Periconia and the family Periconiaceae, the genomes of *Periconia macrospinosa* (Knapp et al., 2018) and *Periconia sp*. R9002 (accession numbers GCA_023627715.1 ASM2362771v1). However, these genome assemblies, made of hundreds of contigs/scaffolds, appear to be much fragmented compared to the genomes that can be obtained nowadays using long read technologies. Therefore, we used a PacBio HiFi sequencing technology to produce highly accurate long reads and assemble the *P. digitata* genome in 13 haploid chromosomes with a total length of ca. 39 Mb. The genome annotation was supported by the assembly of a reference transcriptome made of the transcriptomes of 12 different culture conditions, including stresses, leading to very different mycelium phenotypes. The annotation revealed 15,520 protein-coding genes and conserved InterPro domains could be identified in 60% of the proteins. In addition, we carried out Nano-HPLC-HRMS analyses to characterize the proteome of *P. digitata* and strengthen our functional annotation. The proteomic analysis retrieved and confirmed more than one-third of the predicted proteins no matter the chosen parameter (1 or 2 unique peptides).

Overall, this work generated a high-quality genome completed with rich transcriptomic and proteomic data that will be useful to future research (**Fig.1**). This study would constitute an important resource for our further understanding of the Eukaryotic tree of life and for future comparative genomics (Blaxter et al., 2022). This would help to better delineate the evolution within fungi, a kingdom that is constantly revisited from a systematic point of view on the basis of molecular markers rather than on life history traits. This work will also contribute to bringing new insights into the *Pleosporales*, potentially the largest order of Dothideomycetes that account for more than 300 genera and 4,700 species (Taylor et al., 2015) and contains only 109 sequenced genomes Mycocosm Portals(doe.gov) (2022).

**Fig. 1:**
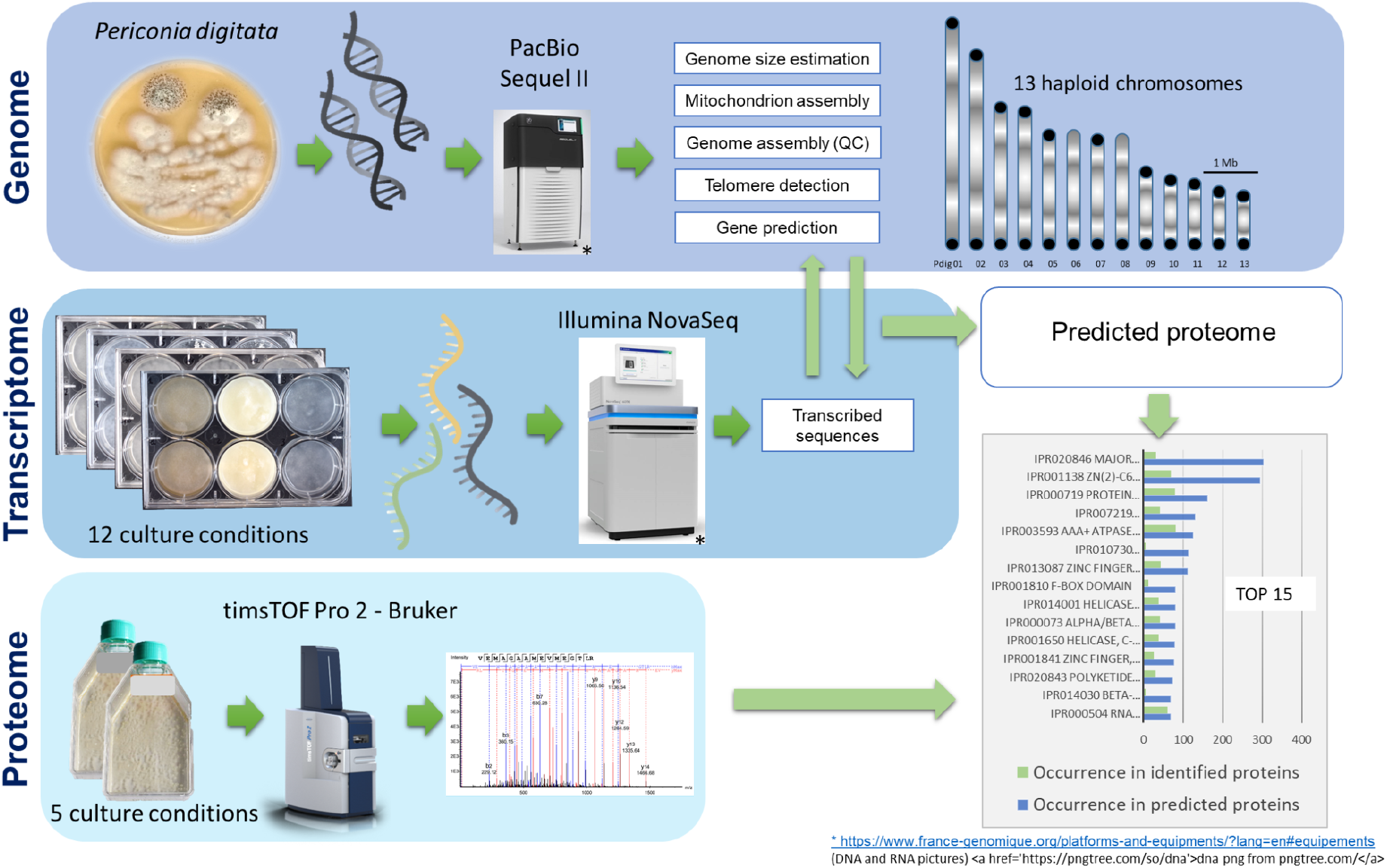
schematic overview of the study design

## Methods

### Strain identification

The strain *Phoma* sp. CNCM I-4278 was previously isolated from the rhizosphere of *Nicotiana tabacum* (cv Xanthi, Solanaceae) grown under controlled conditions (Galiana, Marais, et al., 2011). The fungus was identified according to the closest similarity of its 18S rRNA sequence with those present in GenBank and it was deposited in the National Collection of the Institut Pasteur (CNCM I-4278) (Galiana, Marais, et al., 2011; Galiana, Ponchet, et al., 2011). This first attempt to identify the strain was uncertain since the alignment showed more than 30 mismatches with a putative *Phoma* sp.. Luckily, the increasing availability of fungal sequences in the database allowed a taxonomic reevaluation of the strain.

The fungus was cultivated for 2 weeks on Petri dishes containing Potato Dextrose Agar (PDA - 20 g glucose, 4 g potato extract, 15 g agar, up to 1 L Milli-Q® water) in order to obtain sufficient biomass to perform a DNA extraction. The DNA extraction protocol was optimized in our laboratory starting from previous works (Neuhauser et al., 2011; Plassart et al., 2012; Schneegurt et al., 2003). Briefly, about 100 mg of mycelium were placed in 2 mL Eppendorf tubes with two steel beads and disrupted in a MM301 tissue lyzer (Retsch GmbH, Haan, Germany). Then, a volume of 1 mL of lysis buffer CTAB (28 mM NaCl, 2 mM Tris-base, 0.4 mM Na_2_EDTA, pH 8) plus 2% Polyvinylpyrrolidone (PVP) was added to the samples. The tubes were incubated at 65 °C for 2 h. 400 μL of chloroform:isoamyl alcohol (24:1 v/v) were then added to the samples that were vortexed and centrifuged for 5 min at 13,000 rpm. The supernatant (600 μL) was transferred to a new Eppendorf tube and 100 μL of 10 M ammonium acetate was added; the solution was gently mixed and the samples were incubated at 4 °C for 20 min. The mixture was vortexed and centrifuged for 10 min at 13,000 rpm, the supernatant (650 μL) was transferred to a new Eppendorf tube and one volume of isopropanol (kept at −20 °C) was added. The sample was gently mixed and incubated over night at −20 °C in order to precipitate the DNA. The final pellet was collected by centrifugation for 5 min at 13,000 rpm at 4 °C. The supernatant was discarded and the pellet was washed with 500 μL of 75% aqueous ethanol (kept at −20 °C) and recovered by centrifugation for 2 min at 13,000 rpm. The supernatant was discarded and the pellet was dried under the airflow of a chemical hood. Then, the DNA was resuspended in 30 μL of sterile Milli-Q® water.

The DNA quality and quantity were evaluated using a NanoDrop 2000 (Thermo Scientific, Wilmington, USA). The DNA was stored at −20°C.

The obtained DNA was used to amplify partial sequences of two genetic markers. The primer pairs ITS1/ITS4 (White et al., 1990) and LR0R/LR5 (Vilgalys & Hester, 1990) were used to amplify the internal transcribed spacers and the 28S large ribosomal subunit (nrLSU) region, respectively. The PCR reaction was performed in 25 μL final volumes and consisted of 12.5 μL GoTaq® G2 Hot Start Colorless Master Mix (2X - Promega), 1 μL of each primer (10 μM), 5 μL genomic DNA extract (10 ng/μL) and 5.5 μL nuclease-free water. PCR products were loaded on 2% agarose gel electrophoresis in 0.5X Tris-acetate-EDTA buffer. The gel was stained with ethidium bromide and the PCR products were visualized under UV light. The PCR products were sequenced (accession numbers OP329216 - ITS; and OP329219 - 28S) and aligned with those available in public databases.

This allowed unambiguously identifying the strain *Phoma* sp. CNCM I-4278 as *Periconia digitata* whose name was used further throughout this work.

### *P. digitata* culture conditions

Unless otherwise stated, *P. digitata* was grown in 6-well plates containing 5 mL of sterile media **(Table 1)** that were inoculated with a spore suspension (1.5 x 10^4^ spores/mL final concentration). For the high molecular weight DNA extraction, the fungus was grown on Potato Dextrose Broth - PDB (Potato extract 4 g/L, Dextrose 20 g/L) and incubated at 24 °C, in the dark, for 7 days, prior to DNA extraction.

**Table 1:**
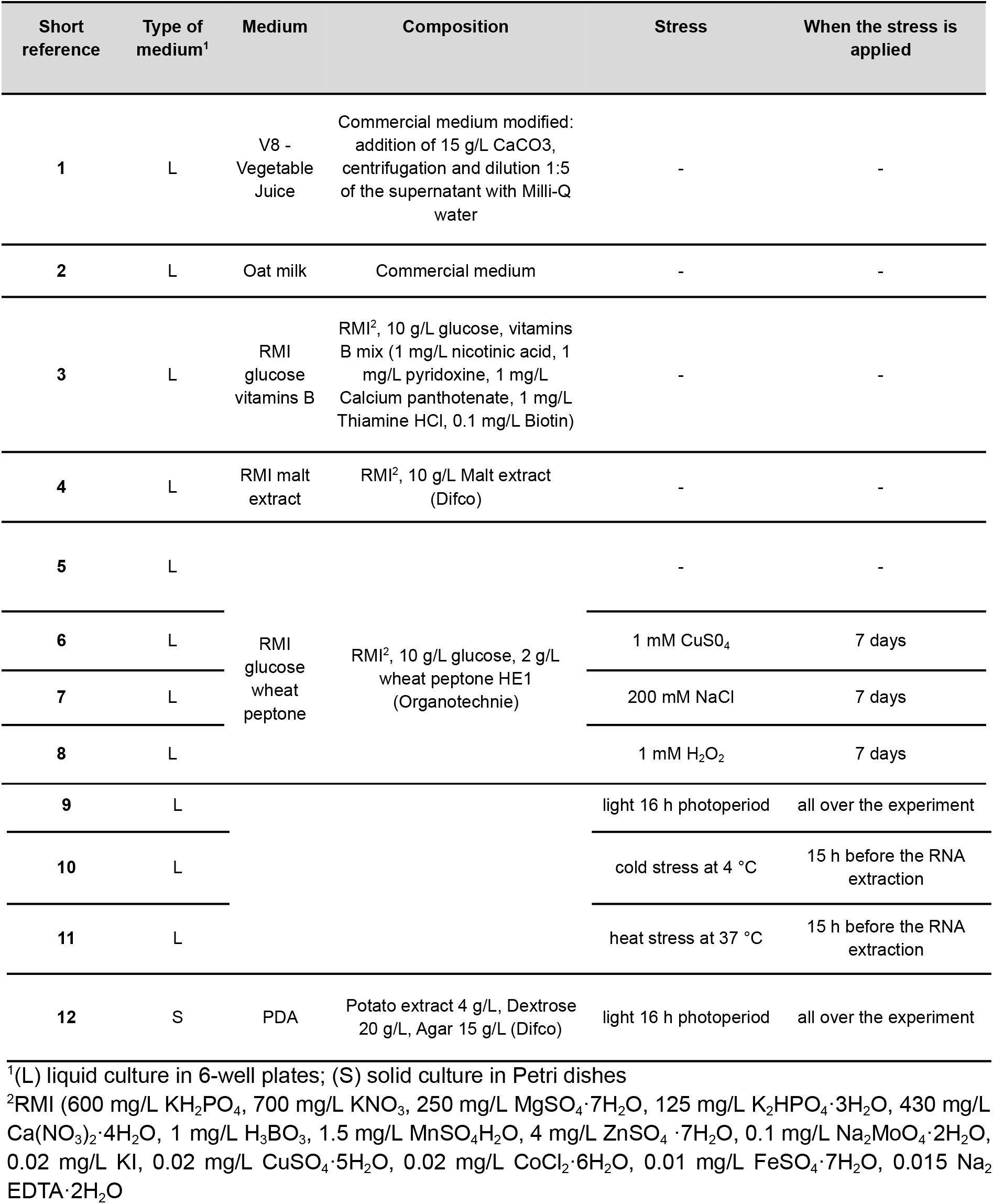
Culture conditions used to obtain a reference transcriptome for *P. digitata*.

In order to generate a reference transcriptome of *P. digitata*, different culture conditions were selected and several stress factors were applied to capture a variety of transcripts and generate an as comprehensive as possible transcriptome. We applied (or not) light, salt, heat, cold, oxidative and heavy metal stresses. We also introduced complex media of plant origin that induced very different phenotypes in terms of mycelium organization. The liquid culture conditions were prepared in 6-well plates with two wells for each condition in the dark. A solid culture was also prepared in Petri dishes (9 cm Ø); the plates were inoculated in three spots with 10 μL of the same spore suspension used for the liquid culture.

The 12 culture conditions, the stress factors and when they were applied are detailed in Table 1. The plates were incubated for 8 days prior to RNA extractions.

### High molecular weight DNA extraction

High molecular weight DNA extraction was performed using the MasterPure™ Complete DNA and RNA Purification Kit from Epicentre. Seven days old-mycelia of *P. digitata* were retrieved from a 6-well plate on PDB as described above and directly ground in liquid nitrogen with sterilized pestles and mortars. The resulting powder was transferred in six microtubes containing 1 μl of proteinase K and 300 μl of tissue and cell lysis solution for each tube and homogenized. All homogenisation steps were performed gently to avoid DNA fragmentation. Tubes were incubated at 65°C for 15 minutes, then cooled down at 37°C before adding 1 μl of 5 μg/μl RNase A and finally incubated for 30 minutes at 37°C. Samples were left on ice for 5 minutes and 175 μl of MPC protein precipitation reagent was added to each sample and mixed. The debris were pelleted by centrifugation at 4°C for 10 minutes at ≥10,000 x g. The supernatant was transferred to a clean microcentrifuge tube. 500 μl of isopropanol was added and tubes were inverted several times before centrifugation at 4°C for 10 minutes. Isopropanol was removed and pellets were rinsed 2 times with 70% ethanol and left dry before solubilization with 35μL of EB buffer (Qiagen). The six samples were pooled together and purified. DNA was analyzed for quality and quantity controls using nanodrop, Qubit and fragment analyser. The final sample used for library preparation contained 19.2 μg of gDNA with an average fragment size of 30,145 bp.

### RNA extraction for transcriptome sequencing

Mycelia from the 12 different culture/stress conditions (Table 1) were retrieved separately and directly ground in liquid nitrogen with sterilized pestles and mortars. The resulting powder for each condition was transferred into 2 to 3 microtubes, depending on the sample quantity. 800 μl of extraction buffer (CTAB 2.5%, PVPP 2%, Tris-HCL 100mM, EDTA 25 mM, NaCl 2M, β-mercaptoethanol 2%) was added to each tube. After 30 minutes of incubation at 65°C, 800 μl of Chloroform/Isoamyl alcohol (CI; v/v; 24/1) was added, and after homogenization, the tubes were centrifuged (16000g - 8 min - 4°C). The supernatant was retrieved and the same volume of water-saturated Phenol (pH 4.5-5) / Chloroform / Isoamyl alcohol (PCI; v/v; 25/24/1) (ca. 700 μl) was added and centrifuged. A second step with CI was carried out and the supernatant was retrieved and mixed with 500 μl NaCl 5M and 500 μl of isopropanol and stored overnight at −20°C. After centrifugation (16000g - 20 min - 4°C), two cleaning steps were carried out with ethanol 70%. The pellet was dried and resuspended in 30 μl of buffer EB (Qiagen). Genomic DNA was removed from samples using the kit TURBO DNA-free (Ambion) following the supplier’s instructions. Samples purity and quality were assessed with a nanodrop and a bioanalyzer. The samples with the best qualitative parameters (1.98<OD_260/280_<2.07; 1.72<OD_260/230_<2.19; 2.5<RIN<5; 3.6 μg<RNA Quantity<15.8 μg) in each condition were kept for library preparation.

### DNA sequencing

Library preparation and sequencing were performed at GeT-PlaGe core facility, INRAE Toulouse according to the manufacturer’s instructions “Procedure & Checklist Preparing HiFi SMRTbell Libraries using SMRTbell Express Template Prep Kit 2.0”. At each step, DNA was quantified using the Qubit dsDNA HS Assay Kit (Life Technologies). DNA purity was tested using the nanodrop (Thermofisher) and size distribution and degradation assessed using the Femto pulse Genomic DNA 165 kb Kit (Agilent). Purification steps were performed using AMPure PB beads (PacBio).15 μg of DNA was purified then sheared at 20 kb using the Megaruptor1 system (Diagenode). Using SMRTbell Express Template prep kit 2.0, a Single strand overhangs removal, a DNA and END damage repair step were performed on 5 μg of sample. Then blunt hairpin adapters were ligated to the library. The library was treated with an exonuclease cocktail to digest unligated DNA fragments. A size selection step using a 9 kb cutoff was performed on the BluePippin Size Selection system (Sage Science) with “0,75% DF Marker S1 High Pass 15-20 kb” protocol. Using Binding kit 2.0 kit and sequencing kit 2.0, the primer V2 annealed and polymerase 2.0 bounded library was sequenced by diffusion loading onto 1 SMRTcell on Sequel2 instrument at 95 pM with a 2 hours pre-extension and a 30 hours movie.

### RNA sequencing

RNAseq was performed at the GeT-PlaGe core facility, INRAE Toulouse. RNA-seq libraries have been prepared according to Illumina’s protocols using the Illumina TruSeq Stranded mRNA sample prep kit to analyze mRNA. Briefly, mRNA were selected using poly-T beads. Then, RNA were fragmented to generate double stranded cDNA and adaptors were ligated to be sequenced. 11 cycles of PCR were applied to amplify libraries. Library quality was assessed using a Fragment Analyser and libraries were quantified by QPCR using the Kapa Library Quantification Kit. RNA-seq experiments have been performed on an Illumina NovaSeq 6000 using a paired-end read length of 2×150 pb with the Illumina NovaSeq 6000 sequencing kits.

### Genome size estimation

We used Jellyfish version 2.2.6 (Marçais & Kingsford, 2011) count and histo commands with a k-mer size of 21 and a maximum multiplicity of 1,000,000 on the PacBio Hi-Fi genome reads to count k-mers and their multiplicity. The output of Jellyfish histo was then used as an input for GenomeScope2 (Ranallo-Benavidez et al., 2020) with a ploidy level of 1 and 2 for genome size and heterozygosity level estimation.

### Mitochondrion assembly

The assembly of the mitochondrial genome sequence was done with ALADIN (https://github.com/GDKO/aladin) using the mitochondrion mode from PacBio HiFi reads. The complete mitochondrial genome of the closest available relative *Phaeosphaeria nodorum* SN15 (NC_009746.1) downloaded from GeneBank was used as a reference seed sequence.

### Genome assembly, QC, Contamination

PacBio Hifi reads with a highly accurate median accuracy of minimum 99.9% (Q30) were used as input for the HiCanu assembler (Nurk et al., 2020). Post-assembly quality control and taxonomic partitioning were assessed with BlobTools (Kumar et al., 2013; Laetsch & Blaxter, 2017). Previously quality-filtered PacBio Hifi reads were mapped back to the assembly with mimimap2 (H. Li, 2018) to estimate contigs coverage. Each contig was assigned a taxonomy affiliation based on BLAST (Altschul et al., 1997; Camacho et al., 2009) results against the NCBI nt database.

### Telomere detection

Terminal telomeric repeats were searched using tidk software v0.1.5 (https://github.com/tolkit/telomeric-identifier). The tidk software explore module was used to search the genome for repeats from length 5 to length 10. Positions of repeats are only reported if they occur sequentially in a higher number than the threshold of 5. The most represented repeat unit was AACCCT with a maximum frequency of 209. Then, this putative telomeric repeat was scanned on the contigs with the tidk search module using a window size of 150 to calculate repeat counts. This information is then used as an input for the tidk plot module to visualize positions of the putative telomeric repeats along each contig sequence.

### Gene prediction

Gene models prediction was done with the fully automated pipeline EuGene-EP version 1.6.5 (Sallet et al., 2019). EuGene has been configured to integrate similarities with known proteins of “ascomycota” section of UniProtKB/Swiss-Prot library (UniProt Consortium 2018), with the prior exclusion of proteins that were similar to those present in RepBase (Bao et al., 2015, p. 201).

The dataset of *Periconia digitata* transcribed sequences generated in this study were aligned on the genome and used by EuGene as transcription evidence. For this, we first assembled *de novo* using Trinity (Haas et al., 2013) the transcriptomes of *P. digitata* obtained from the twelve above-described conditions and for a given trinity locus we only retained the transcript returning the longest ORF. Finally, only *de novo* assembled transcripts that aligned on the genome on at least 30% of their length with at least 97% identity were retained.

The EuGene default configuration was edited to set the “preserve” parameter to 1 for all datasets, the “gmap_intron_filter” parameter to 1 and the minimum intron length to 35 bp. Finally, the Fungi specific Weight Array Method matrices were used to score the splice sites (available at this URL: http://eugene.toulouse.inra.fr/Downloads/WAM_fungi_20180126.tar.gz).

### Genome and protein set completeness assessment

We used BUSCO (Manni et al., 2021) version 5.2.2 in protein and genome modes with the eukaryota odb10 dataset of 255 BUSCO groups and the fungi odb10 dataset of 758 BUSCO groups to assess the completeness of the predicted protein set as well as the genome assembly. We compared BUSCO scores to those obtained for the *Periconia macrospinosa* genome and predicted proteins (Knapp et al., 2018).

### Functional annotation

All predicted proteins were scanned for the presence of conserved protein domains and motifs using InterProScan v.5.51-85.0 (Jones et al., 2014) with the options -iprlookup, -goterms and -pa to assign Gene Ontology (GO) terms, MetaCyc and Reactome biochemical pathways based on detection of Interpro domains.

### Gene prediction and functional annotation of mitochondrion

The annotation was performed using MITOS2 (Donath et al., 2019) including the ncRNA (t- and r-RNA) and the protein coding sequences with the table 4 codon usage. The gene predictions were refined using the assembled Trinity transcripts aligned on the mitogenome by direct translation in ORFfinder as well as annotation with smartBlast (https://www.ncbi.nlm.nih.gov/orffinder/), and intron reconstruction with BioEdit v 7.0.5.3 (Hall, 1999). We also compared our results to the following Pleosporales available mitogenomes: NC_058694 (*Edenia gomezpompae*, 37 kb, 14 ORF), NC_040008 (*Coniothyrium glycines*, 98 kb, 35 ORF), NC_026869 (*Shiraia bambusicola*, 39 kb, 17 ORF) and NC_035636 (*Pithomyces chartarum*, 69 kb, 37 ORF) in addition to this of *Phaeosphaeria nodorum* (see mitochondrion assembly). The concatenated file obtained from MITOS2 and protein-coding sequences coordinates was used for genbank submission (OP787475) and mitogenome drawing by OGDRAW version 1.3.1 (Greiner et al., 2019).

### Protein extraction and sample preparation for proteomics analysis

*P. digitata* was cultivated, in 500 mL plastic Roux bottles, in 50 mL of five different sterile liquid media (RMI free of asparagine and vitamins, RMI supplemented with B vitamins, RMI plus wheat peptone, RMI plus *Citrus* pectin, RMI plus Guar gum, RMI plus malt) over 7 days in the dark at 24 °C. The media were chosen for their ability to change the strain phenotype (see RNA extraction above); two biological replicates were prepared. The mycelium was recovered by filtration on GF/C Whatman glass filter and rinsed with water. It was ground using liquid nitrogen then sequentially extracted using successive buffers starting with a Tris buffer 20 mM pH 8, 10mM DTT (dithiothreitol),then the same buffer supplemented with 200 mM NaCl buffer, the same buffer supplemented with 8 M urea and finally the same buffer with 6 M guanidine hydrochloride (1 mL per 300 mg fresh material weight). Centrifugations (13,200 rpm, 5 min, 4°C) were achieved after each extraction and the 4 successive supernatants were recovered. The two first buffers (alone and saline) were immediately adjusted to 8 M urea. All these fractions were incubated at 37 °C for 15 min then alkylated by iodoacetamide (41.6 mM) during 15 minutes at room temperature. After two buffer exchanges with trypsin buffer on Vivaspin 15R (5 kDa) columns (Sartorius), the samples were digested by trypsin with 1/80 (w/w) trypsin/total protein ratio (Sequencing Grade Modified Trypsin, Promega) according to the manufacturer recommendations. The residual pellets after successive extractions were suspended in 1 mL of Tris 20 mM pH 8, 10 mM DTT, 8M urea then incubated at 37 °C for 15 min and alkylated with 41.6 mM of iodoacetamide for 15 min at room temperature with gentle mixing. After centrifugation (see above), the pellets were rinsed/centrifuged 2 fold in the trypsin buffer. The final pellet suspension was directly digested by trypsin (1 μg/pellet) overnight under gentle agitation on a rotator mixer. After centrifugation, the supernatants were recovered. The samples were then cleaned-up on a C18 SPE cartridge (100 mg, NUCLEODUR® 100-30 C18 end-capped, Macherey Nagel) equilibrated in H2O 0.1% formic acid (FA). After washing with H2O 0.1% FA, peptides were eluted with acetonitrile (CH3CN)-0.1% FA / water-0.1% FA 40/60 and 70/30 (v/v). Both fractions were pooled.

### Proteomics analysis

The samples were analyzed by nanoUHPLC-HRMS (nanoElute – timsTOF Pro, Bruker Daltonics). 5 μL of sample were injected on an Aurora column (75μm id x 250 mm, C18, 1.6 μm, ionOpticks) with a flow rate of 200 nL/min at 50°C. The mobile phase was a gradient of CH3CN-0.1 % FA (B) in 0.1% FA-H_2_O (A) as follows: 5% B for 1 min, 5% to 13% of B for 18 min, 13% to 19% of B for 7 min, 19% to 22% of B for 4 min, 22% to 95% of B for 3 min.

The timsTOF Prowas equipped with the CaptiveSpray nano-electrospray ion source. MS and MSMS data were acquired in a positive mode, in a PASEF (Parallel Accumulation -Serial Fragmentation) data dependent acquisition (DDA), TIMS ON mode from 100 to 1700 m/z mass range (TimsControl version 2.0.53.0). Ion mobility resolution (1/K0) was set to 0.70–1.10 V.s/cm^2^ over a ramp time of 180 ms. To exclude low m/z, singly charged ions from PASEF precursor selection, a polygon filter was applied in the m/z and ion mobility space.

Analyses were processed by Peaks Studio X Pro (version 10.6, bioinformatics Solutions Inc.) MSMS raw data were processed using Peaks solution applying 3 levels of identification. MSMS spectra were matched against the *P. digitata* Y3 predicted proteome (merged core and mitochondrial genomes), including the MaxQuantcontaminant database (contaminants.fasta, MaxQuant 2.1). Parameters were set as follows: protein FDR<1%, decoy fusion method, cysteine carbamidomethylation as a fixed modification, 2 miscleavage, 3 to 5 post-translational modifications (PTM) per peptide. The MSMS spectra that did not match with these parameters were processed again, looking for other possible PTM and amino acid mutations to enrich the list of identified proteins. The obtained results were merged in a list of significant proteins (supplementary data).

## Data records

ITS and 28S sequences that have been used to identify *P. digitata* are available in NCBI (accession numbers OP329216 and OP329219, respectively).

All Illumina and PacBio HiFi raw data used to assemble the genome and transcriptomes are publicly available at the EMBL-EBI’s European Nucleotide Archive under the project number **PRJEB55037** (https://www.ebi.ac.uk/ena/browser/view/PRJEB55037). The detailed list of raw data accession numbers are presented in the table below.

|  | Run Accession | Sample | Experiment |
| --- | --- | --- | --- |
| RNA of *P. digitata* condition 12 | ERR10025227 | ERS12553875 | ERX9565917 |
| RNA of *P. digitata* condition 11 | ERR10025174 | ERS12553874 | ERX9565864 |
| RNA of *P. digitata* condition 10 | ERR10025118 | ERS12553873 | ERX9565808 |
| RNA of *P. digitata* condition 9 | ERR10025114 | ERS12553872 | ERX9565804 |
| RNA of *P. digitata* condition 8 | ERR10025101 | ERS12553871 | ERX9565791 |
| RNA of *P. digitata* condition 7 | ERR10025086 | ERS12553870 | ERX9565776 |
| RNA of *P. digitata* condition 6 | ERR10025067 | ERS12553869 | ERX9565757 |
| RNA of *P. digitata* condition 5 | ERR10025065 | ERS12553868 | ERX9565755 |
| RNA of *P. digitata* condition 4 | ERR10025044 | ERS12553867 | ERX9565734 |
| RNA of *P. digitata* condition 3 | ERR10024935 | ERS12553866 | ERX9565625 |
| RNA of *P. digitata* condition 2 | ERR10024912 | ERS12553865 | ERX9565602 |
| RNA of *P. digitata* condition 1 | ERR10020517 | ERS12553864 | ERX9561207 |
| DNA of *P. digitata* | ERR10009562 | ERS12521256 | ERX9550522 |

The mass spectrometry proteomic raw data are available in the ProteomeXchange Consortium via the PRIDE [1] partner repository with the dataset identifier **PXD038112** and **PXD038175**.

All the analyzed data are publicly available at https://entrepot.recherche.data.gouv.fr/dataverse/pdig. The detailed list of analyzed files are presented in the table below.

| doi accession (https://doi.org/) | Description |
| --- | --- |
| 10.57745/FB9OVN | genome assembly fasta file |
| 10.57745/BAPYY5 | gene models prediction and eugene statistics files |
| 10.57745/5ODV0C | genes fasta file |
| 10.57745/71DS6F | CDS fasta file |
| 10.57745/TV4EB7 | predicted proteins fasta file |
| 10.57745/GU67TE | functional annotation provided by interproscan |
| 10.57745/FHNYJU | proteins identified by mass spectrometry. It includes the list of proteins identified with more than 1 peptide, only 1 peptide, the Post Translational Modifications, proteins displaying a signal peptide. |

The complete mitochondrial genome assembly of *P. digitata* can be retrieved at the NCBI through the accession number **OP787475** (https://www.ncbi.nlm.nih.gov/search/all/?term=OP787475)

## Technical validation

### Contamination assessment

Blobtools analysis showed that all the contigs formed a dense blob at a homogenous coverage (550X) and GC content (49%) indicating no evidence for contamination (i.e. no contig deviates from this distribution) **(Fig. 2)**. Moreover, the taxonomic affiliation analysis based on homology, using BLAST against the NCBI’s nt library, showed that all contigs are of Ascomycota origin, which is consistent with the absence of evident contamination **(Fig. 3)**.

**Fig. 2:**
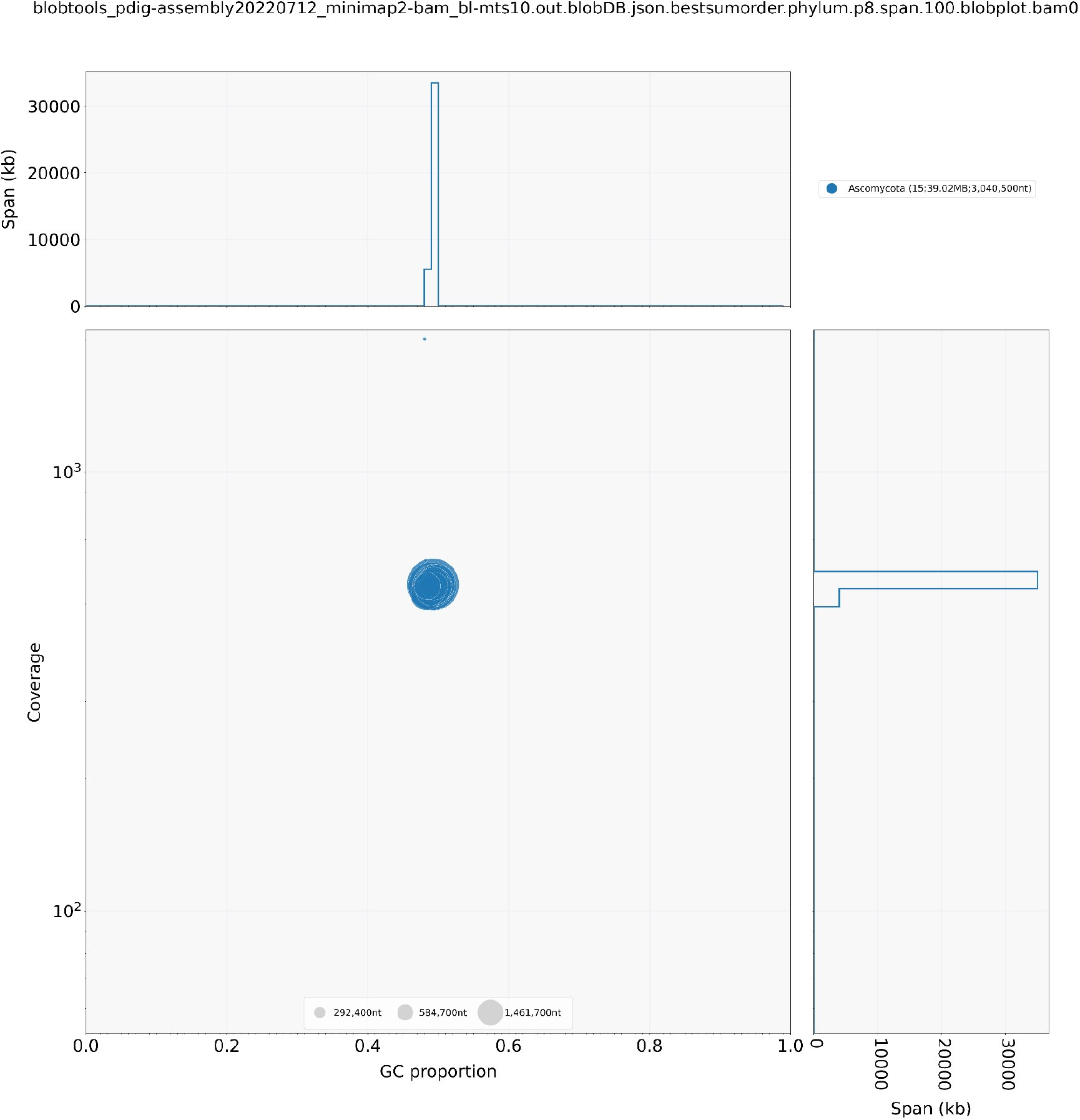
BlobPlot of the genome assembly. Each circle is a contig proportionally scaled by contig length and coloured by taxonomic annotation based on BLAST similarity search results. Contigs are positioned based on the GC content (X-axis) and the coverage of PacBio reads (Y-axis).

**Fig. 3:**
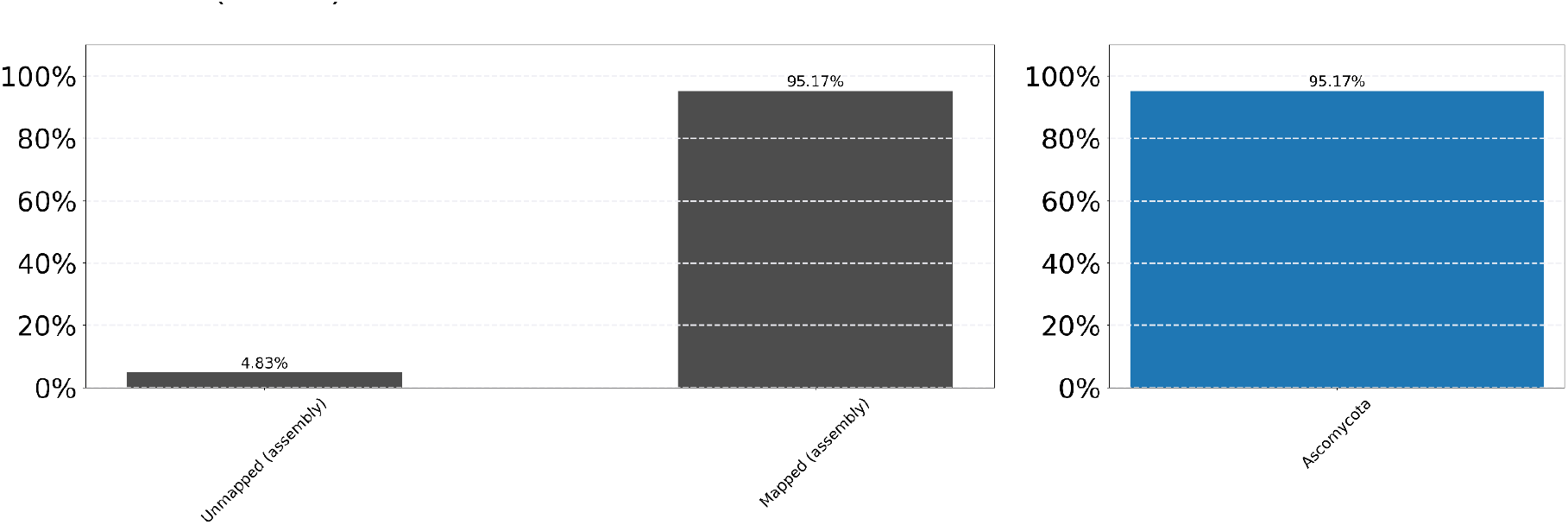
ReadCovPlot. Mapped reads are shown by the taxonomic group at the rank of ‘phylum’.

### Genome size estimation and *de novo* assembly

Based on k-mer multiplicity distribution using GenomeScope2, both 1n and 2n models converged in showing one single peak at a very high coverage of ca. 560X. The GenomeScope model fit values were slightly higher for the haploid (1n) model (92.59% - 94.61%) than for the diploid (2n) model (92.59% - 94.31%). Collectively, these results strongly suggest a haploid genome sequenced at a very high coverage. The haploid 1n model returned an estimated genome size of ca. 36 Mb with an error rate of ca. 0.43% (**Fig. 4).**

**Fig. 4:**
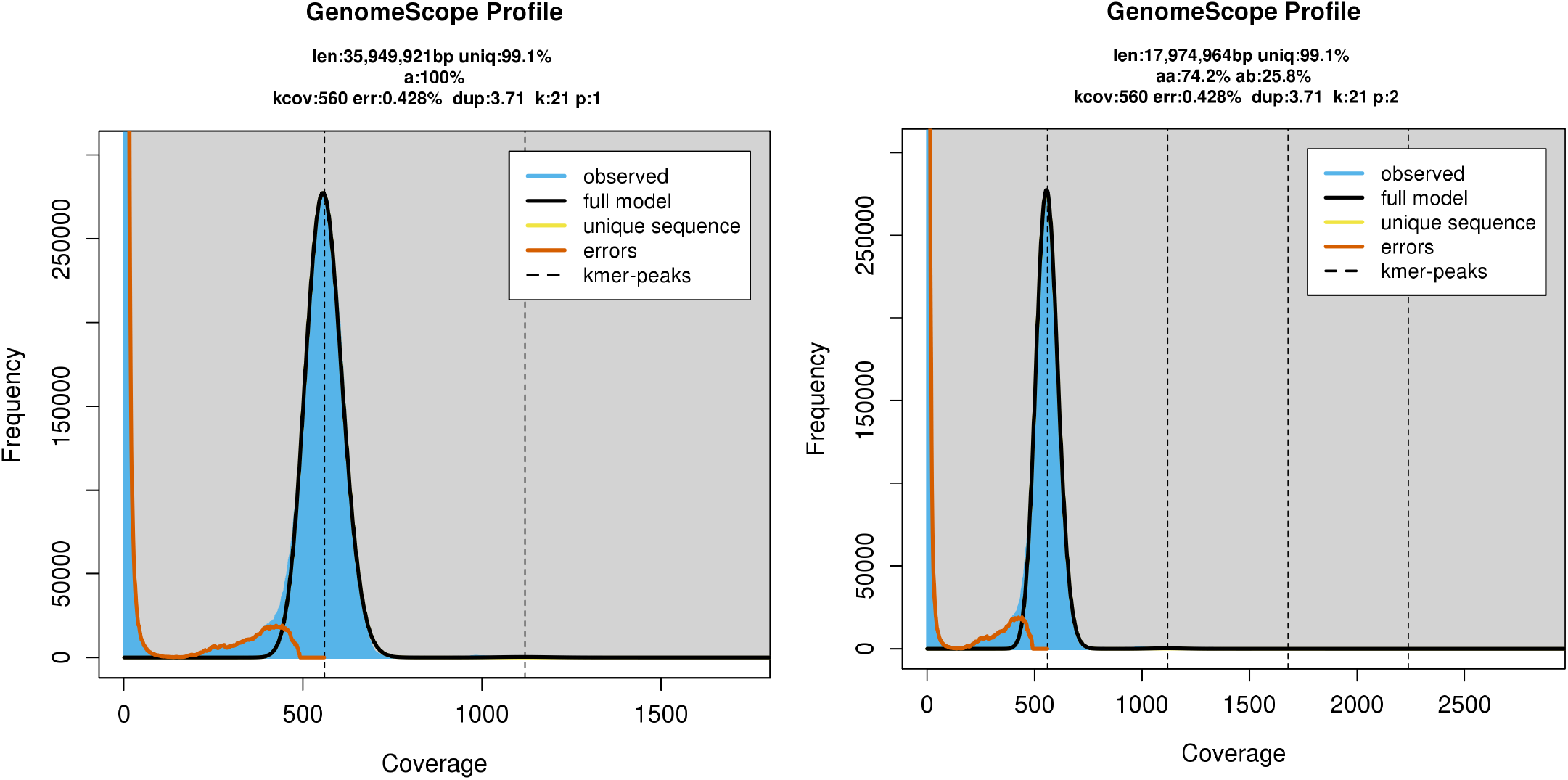
GenomeScope2 k-mer profiles for the haploid (1n) (left) and diploid (2n) (right) models. Coverage (kcov), error rate (err.), haploid genome size estimation (len.), k-mer size (k), ploidy level (p)

The HiCanu assembler yielded a genome assembly that was ca. 39 Mb long, consistent with haploid genome size estimated with k-mers. The genome was assembled in 15 contigs with a N50 value of 3 Mb and a L50 of 5 (i.e. half of the genome is present in the 5 biggest contigs) (**Table 2**). Among the 15 contigs, HiCanu generated two outliers of 27-28kb (Pdig14 and Pdig15, **Table 3**). Both contained only rDNA repeats that can be partially aligned at the end of Pdig08, which exclusively exhibits rDNA repeats.

**Table 2:**
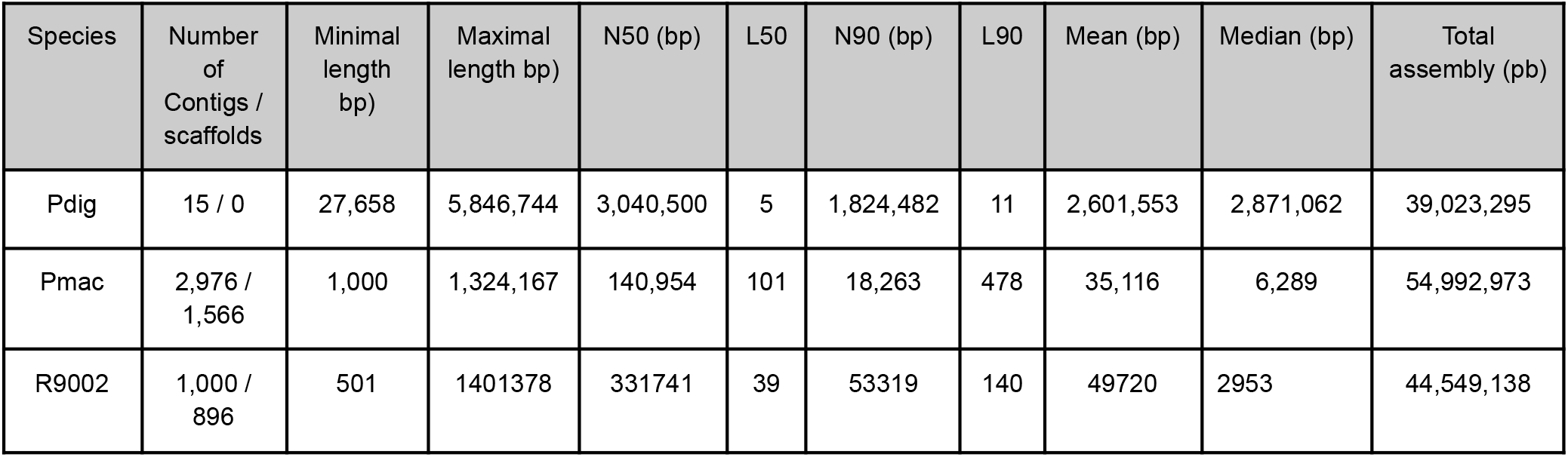
Metrics of *Periconia digitata’s* genome assembly (Pdig) and comparison to *P. macrospinosa* (Pmac) (assembly GCA_003073855.1 Perma1, Knapp et al. 2018) and *P*. R9002 (assembly GCA_023627715.1 ASM2362771v1)

**Table 3:** Metrics of *Periconia digitata’s* contigs and telomere detection

| ContigsID | length (bp) | GC% | Telomere detection (Tidk) |
| --- | --- | --- | --- |
| Pdig01 | 5,846,744 | 49.26 | both ends |
| Pdig02 | 4,988,348 | 49.15 | both ends |
| Pdig03 | 3,687,698 | 49.32 | both ends |
| Pdig04 | 3,596,248 | 49.34 | both ends |
| Pdig05 | 3,040,500 | 49.32 | both ends |
| Pdig06 | 3,027,279 | 49.72 | start |
| Pdig07 | 2,891,359 | 49.35 | both ends |
| Pdig08 | 2,871,062 | 49.26 | start |
| Pdig09 | 2,118,696 | 48.31 | both ends |
| Pdig10 | 1,885,720 | 49.35 | both ends |
| Pdig11 | 1,824,482 | 48.93 | both ends |
| Pdig12 | 1,658,798 | 49.42 | both ends |
| Pdig13 | 1,530,560 | 48.47 | both ends |
| Pdig14 | 28,143 | 48.07 | none |
| Pdig15 | 27,658 | 48.21 | none |

**Table 4:**
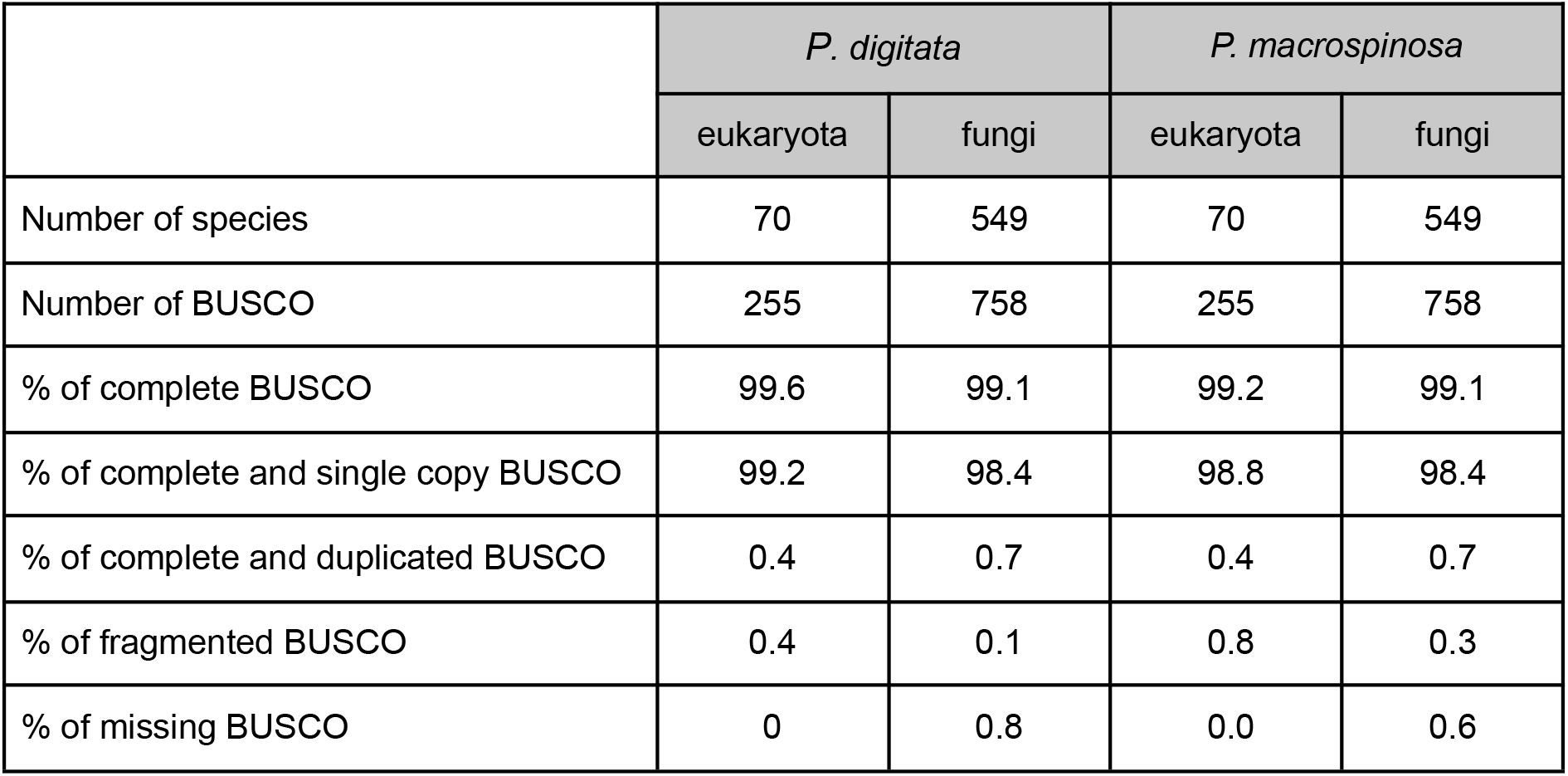
BUSCO scores for the genome of *P. digitata* and *P. macrospinosa* using the eukaryota odb10 and the fungi odb10 datasets

|  | *P. digitata* |  | *P. macrospinosa* |  |
| --- | --- | --- | --- | --- |
|  | eukaryota | fungi | eukaryota | fungi |
| Number of species | 70 | 549 | 70 | 549 |
| Number of BUSCO | 255 | 758 | 255 | 758 |
| % of complete BUSCO | 99.6 | 99.1 | 99.2 | 99.1 |
| % of complete and single copy BUSCO | 99.2 | 98.4 | 98.8 | 98.4 |
| % of complete and duplicated BUSCO | 0.4 | 0.7 | 0.4 | 0.7 |
| % of fragmented BUSCO | 0.4 | 0.1 | 0.8 | 0.3 |
| % of missing BUSCO | 0 | 0.8 | 0.0 | 0.6 |

The repeat sequence (AACCCT)n we have identified at the terminal regions of the contigs corresponds to the reverse complement of the (TTAGGG)n telomeric repeat widely conserved in vertebrates, many other animals, plants as well as several different eukaryotes, including fungal species.

It is worthy to note that the telomeric repeats were detected at both ends of 11 out of 13 contigs. Telomeric repeats were detected at only one end in the contigs Pdig06 and Pdig08 (**Table 3**). Overall, we obtained a highly-contiguous genome assembly for *Periconia digitata* that is structured in 13 chromosomes in its haploid mycelium with a total length of 38,967,494 bp.

By comparison with the two other genomic resources available for the *Periconia* genus (P. *macrospinosa* and *P. sp*. R9002), the assembly of *P. digitata* greatly improves the resolution of Periconia’s genome structure. *P. macrospinosa* (assembly GCA_003073855.1 Perma1, Knapp et al. 2018) and *P. sp*. R9002 (assembly GCA_023627715.1 ASM2362771v1) assemblies (54.99 Mb and 44.55 Mb, respectively) are made of 2,976 / 1,566 and 1,000 / 896 contigs / scaffolds, with a N50 of 140,954 bp and 237,696 bp, respectively (Table 2).

### Genome completeness assessment

BUSCO (v5.2.2) analysis at the genome level indicated that 99.6% and 99.1% of nearly-universal single copy genes from the eukaryota and fungi datasets, respectively, were retrieved in full-length. Only 0.4 and 0.7% of eukaryotic and fungal BUSCO genes were identified as duplicated, consistent with a genome assembled in a haploid state with no evidence for substantial gene duplications **(Table 4)**. Although the genome of *P. macrospinosa* was considerably more fragmented, the gene content completeness was comparable to that of P. digitata according to BUSCO metrics.

### Genome annotation and assessment of predicted proteins

Using the Eugene-EP pipeline, 15,815 genes were predicted including 15,520 protein-coding genes and 295 non-coding genes. Genes cover 26.3 Mb (~68%) of the genome assembly. The coding portion accounts for 51% of the assembly and spliceosomal introns were detected in 71% of protein coding-genes, with an average of 2.61 exons per gene.

A BUSCO analysis in protein mode, revealed that 96.9% and 97.3 % of complete eukaryotic and fungal BUSCO proteins, respectively, were found in the predicted proteome dataset **(Table 5)**. The metrics obtained on *P. macrospinosa* annotation were very similar, suggesting again a complete gene set despite a fragmented genome assembly.

**Table 5:** BUSCO scores for the proteins of *P. digitata* and *P. macrospinosa* using the eukaryota odb10 and the fungi odb10 datasets

|  | *P. digitata* |  | *P. macrospinosa* |  |
| --- | --- | --- | --- | --- |
|  | eukaryota | fungi | eukaryota | fungi |
| Number of proteomes | 70 | 549 | 70 | 549 |
| Number of BUSCO | 255 | 758 | 255 | 758 |
| % of complete BUSCO | 96.9 | 97.3 | 97.6 | 98.5 |
| % of complete and single copy BUSCO | 96.9 | 96.8 | 97.6 | 97.8 |
| % of complete and duplicated BUSCO | 0.0 | 0.5 | 0.0 | 0.7 |
| % of fragmented BUSCO | 1.2 | 0.8 | 1.2 | 0.4 |
| % of missing BUSCO | 1.9 | 1.9 | 1.2 | 1.1 |

### Functional annotation

Conserved InterPro domains and motifs were identified on 61.1% of the 15,520 predicted protein sequences. The 9,479 annotated proteins returned 7,713 different InterPro domains.

The top 15 interpro homologous superfamilies and domains contained some large gene families found in most organisms together with domains restricted to fungi. Among the detected domains, we noticed the presence of the HET (Heterokaryon incompatibility) domain (**Fig. 5**) which is specific to Ascomycota (Espagne et al., 2002) and domains associated with key enzymes for many secondary metabolites biosynthesis in fungi such as the beta-ketoacyl synthase or the polyketide synthase, enoylreductase domains ((Nicolaisen et al., 1997).

**Fig. 5:**
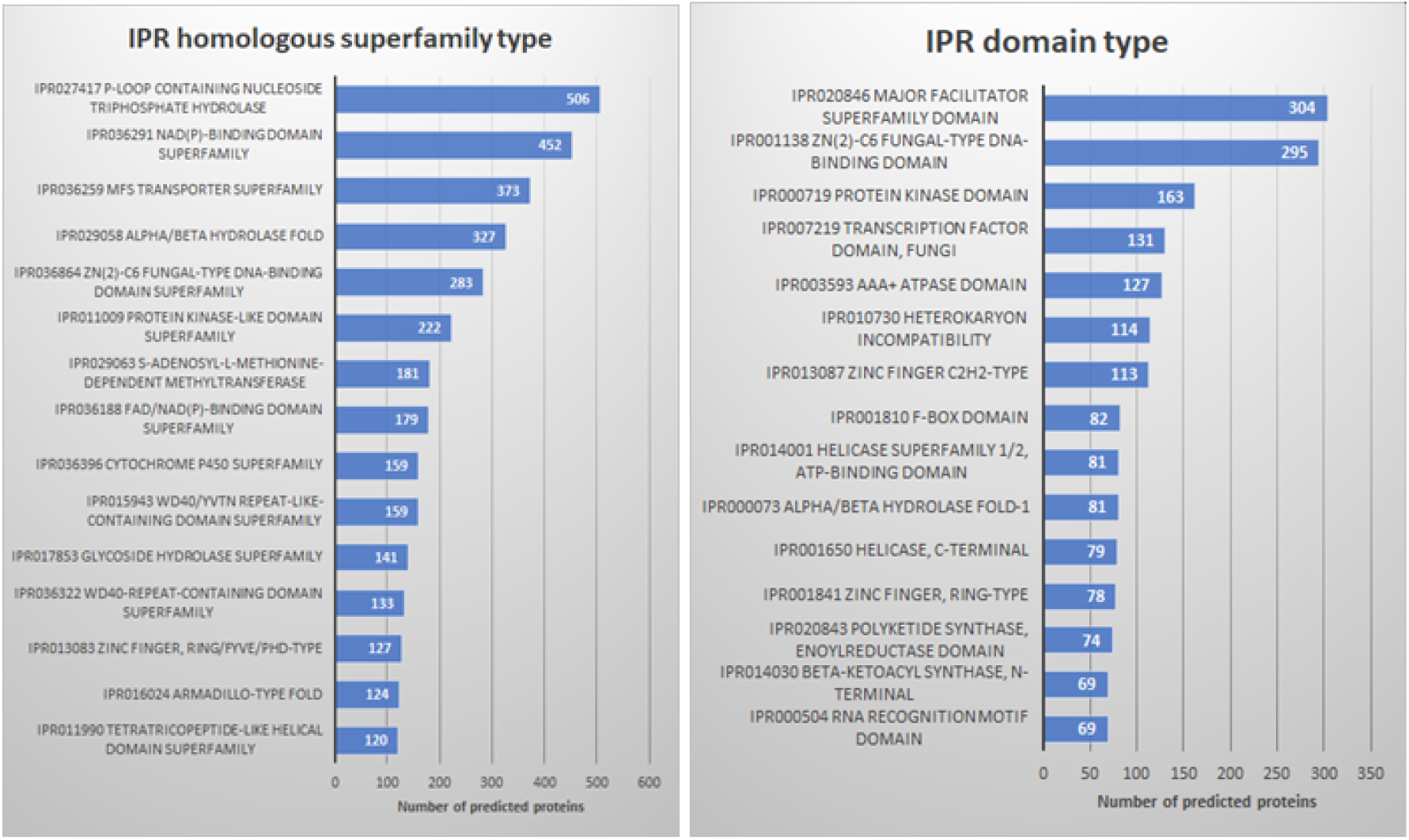
Interpro functional annotation of the *P. digitata* predicted proteome. The Top 15 homologous superfamilies (a) and domains (b) are indicated.

Using SinalP (v6.0) (Teufel et al., 2022), we assessed the number of predicted proteins that contained a putative signal peptide for secretion. Among the 15,520 proteins, 1,597 (10.3%) were predicted to display a signal peptide thus potentially to be addressed to extracellular space or the membrane. This value is in the upper section of the range observed in fungal proteomes, e.g. 8.5% in *Trichoderma asperellum* (Zheng et al., 2022), 1.1% - 12% in 132 Zygomycota proteomes (Chang et al., 2022), 3% - 10% obtained in 49 fungal proteomes (Pellegrin et al., 2015).

### Proteomics support of predicted proteins

Of the 15,551 predicted proteins, 6,598 (42.4%) returned matches with at least 2 unique peptides (−10lgP >50) at a maximum FDR of 1%.The identification reached 46.9 % after inclusion of proteins identified with a single peptide (698, −10lgP>50). This value is in line with the numbers observed among the top 15 interpro homologous superfamilies and domains identified (**Fig. 6).** Although 6,041 proteins (38.9% of the predicted ones) returned no InterPro annotation, 883 of them (14.6%) were identified by proteomics.

**Fig. 6:**
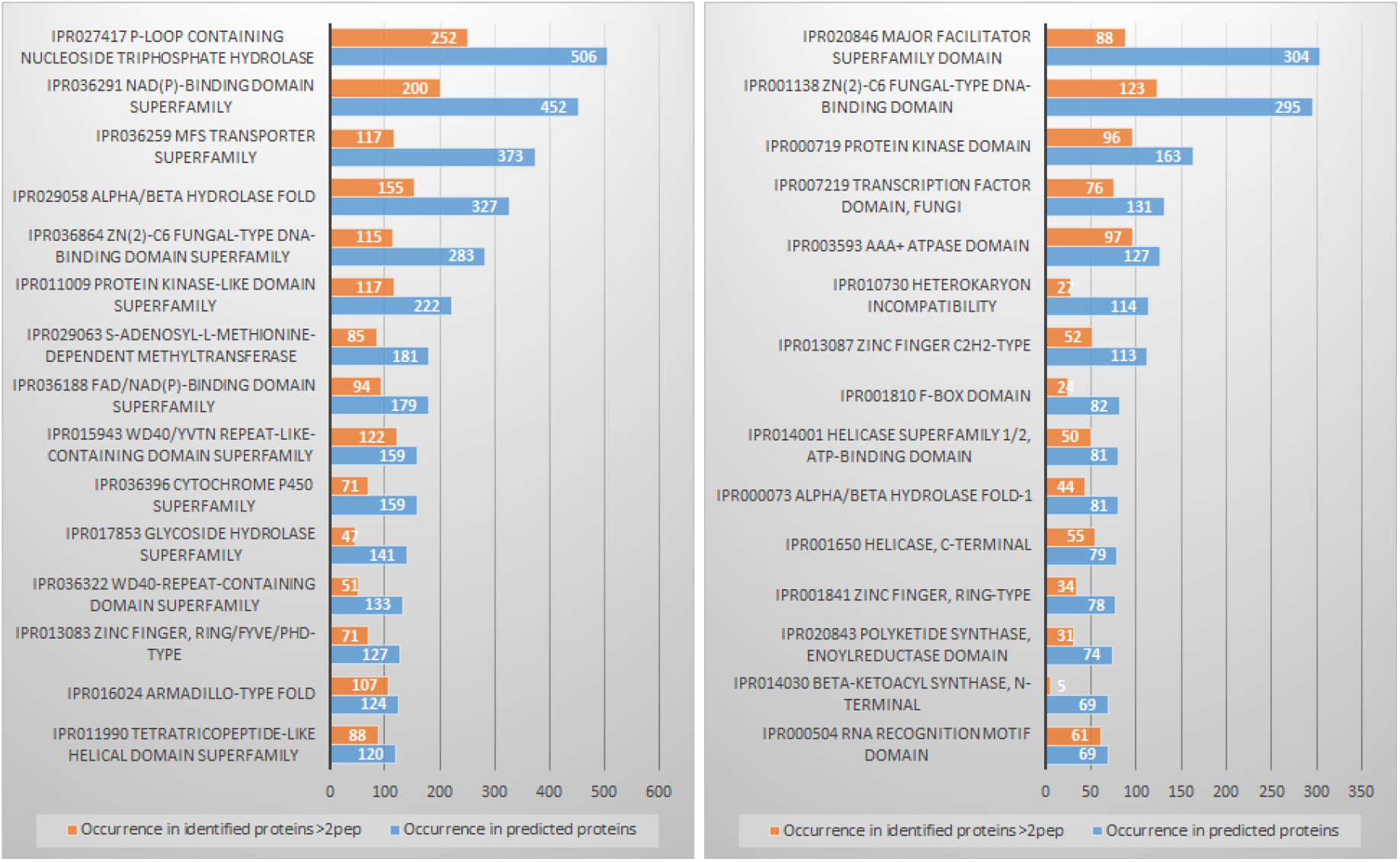
Top 15 homologous superfamilies (a) and domains (b) obtained from Interpro predictions (blue) together with the corresponding homologous superfamilies and domains in proteins identified by proteomics (orange).

**Figure 6:**
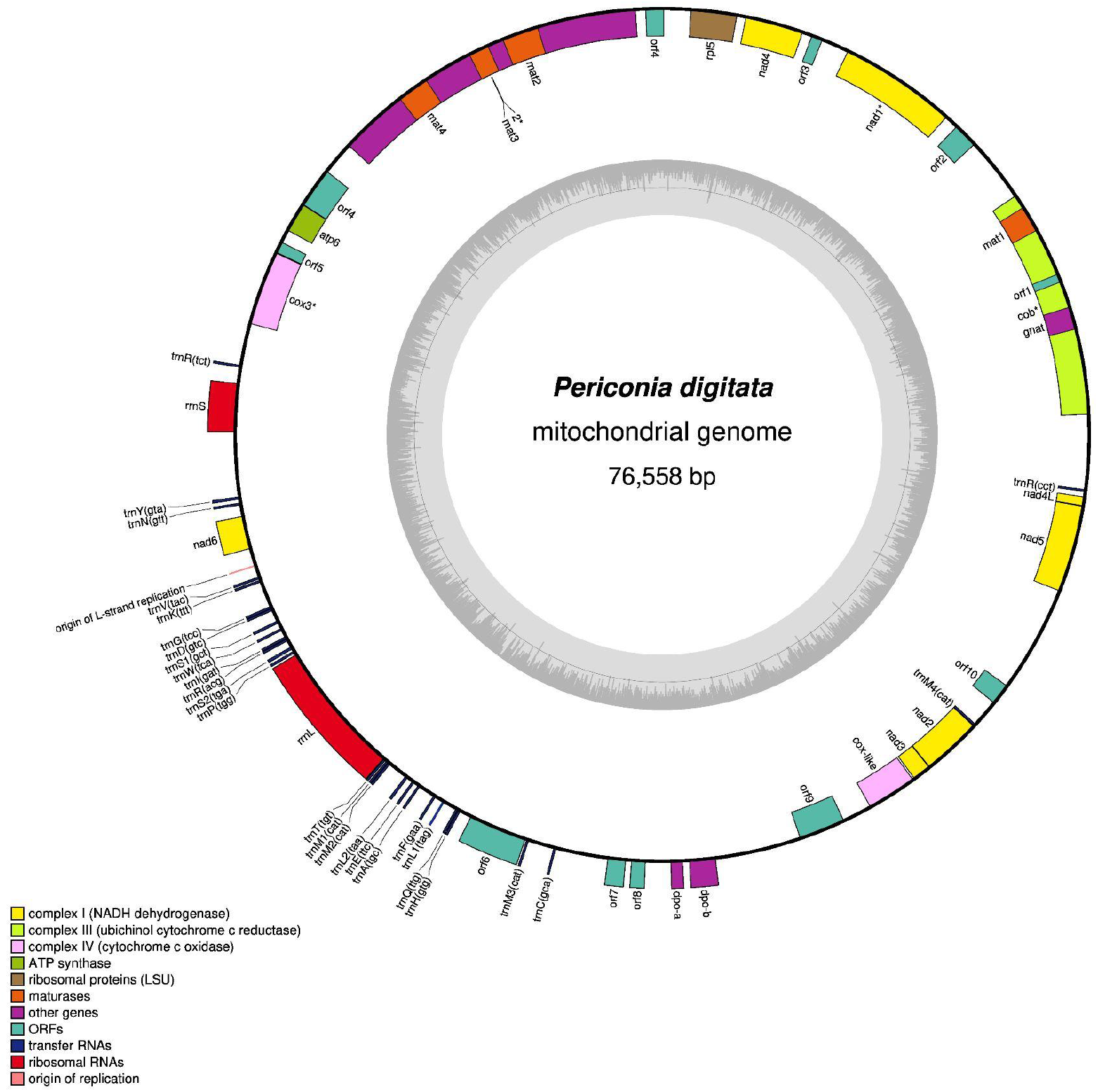
Mitochondrial genome of *Periconia digitata*

In addition, many post translational modifications (PTM, A score >20) were detected (**Table 6**). Phosphorylation mainly concerned proteins predicted to be involved in transport (13), ubiquitin trafficking (13) cytoskeleton (12) and transcription (8). Acetylation and methylation mainly concerned proteins predicted to interact with nucleic acids (from histones to ribosomal proteins) or to have diverse enzymatic activities.

**Table 6 :**
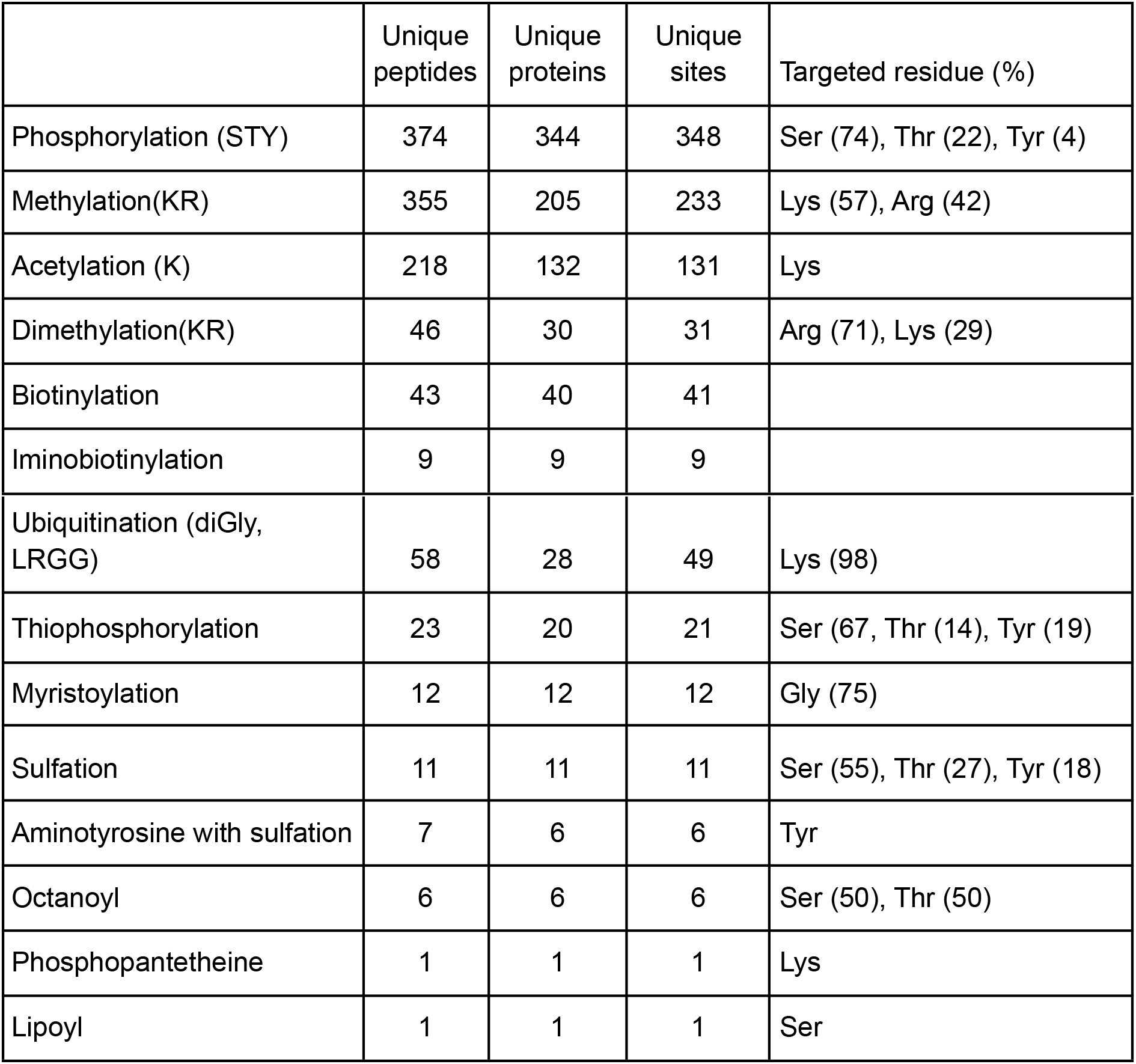
Detected post translational modifications (PTM) with a A score >20 (equivalent to pvalue < 0.01)

### Mitochondrial genome assembly and annotation

The assembled mitochondrial genome was 76,558 bp long with 27.56% of GC (**Figure 6**). As for other Pleosporales mitogenomes, we retrieved a complete set of tRNA, among which some were in multiple copies(Met, Ser, Leu) as well as rnl and rns. Out of the 31 predicted proteins encoded in the mitochondrion, 14 were identified by proteomics including typical enzymes, one ribosomal protein and 3 hypothetical proteins (**Table 7**).

**Table 7:** list of the 14 mitochondrial proteins identified by proteomics

| Accession | Coverage (%) | Unique peptides | Annotation |
| --- | --- | --- | --- |
| PdigMGS11 | 34 | 72 | cox1&2 |
| PdigMGS09 | 47 | 24 | rpl5 |
| PdigMGS30 | 21 | 22 | nad5 |
| PdigMGS25 | 36 | 21 | hypothetical protein |
| PdigMGS15 | 35 | 15 | hypothetical protein |
| PdigMGS28 | 15 | 10 | nad2 |
| PdigMGS08 | 15 | 9 | nad4 |
| PdigMGS27 | 31 | 8 | nad3 |
| PdigMGS29 | 31 | 8 | hypothetical protein |
| PdigMGS18 | 6 | 6 | cox3 |
| PdigMGS01 | 12 | 6 | cob |
| PdigMGS16 | 15 | 5 | atp6 |
| PdigMGS06 | 4 | 2 | nad1 |
| PdigMGS31 | 9 | 1 | nad4L |

## Acknowledgments

This work was funded by the French Public Bank of Investment (BPI France) within the framework of the PSPC Project “SOLSTICE” (SOLutionS pour des Traitements Intégrés dans une Conduite Environnementale).

The authors thank the “Plant Health and Environment” INRAE department for its constant support. We are grateful to the genotoul bioinformatics platform Toulouse Occitanie (Bioinfo Genotoul, https://doi.org/10.15454/1.5572369328961167E12) for computing resources. We are grateful to the bioinformatics and genomics platform, BIG, Sophia Antipolis (ISC plantBIOs, https://doi.org/10.15454/qyey-ar89) for computing and storage resources.

## Author contributions

M.P., E.B., C.R. C.V.G and E.G.J.D conceived the study. C.R., EB, M.P., E.G.J.D and C.V.G. wrote the manuscript with input from all authors. C.R., C.K. and E.G.J.D designed and carried out the bioinformatic analyses and data transfer. E.B. produced the biological material. C.V.G. carried out the nucleic acid extractions. M.P., A.S. and M.M. designed and carried out the proteomic analyses. M.G. performed the PacBio HiFi library and sequencing. A.L performed the RNA libraries preparations and Illumina sequencing.

## References

Altschul, S. F., Madden, T. L., Schaffer, A. A., Zhang, J., Zhang, Z., Miller, W., & Lipman, D. J. (1997). Gapped BLAST and PSI-BLAST: a new generation of protein database search programs. Nucleic Acids Res, 25(17), 3389–3402.

Azhari, A., & Supratman, U. (2021). The Chemistry and Pharmacology of Fungal Genus Periconia: A Review. Scientia Pharmaceutica, 89(3), 34. https://doi.org/10.3390/scipharm89030034

Bao, W., Kojima, K. K., & Kohany, O. (2015). Repbase Update, a database of repetitive elements in eukaryotic genomes. Mobile DNA, 5(1), 11. https://doi.org/10.1186/s13100-015-0041-9

Blaxter, M., Archibald, J. M., Childers, A. K., Coddington, J. A., Crandall, K. A., Di Palma, F., Durbin, R., Edwards, S. V., Graves, J. A. M., Hackett, K. J., Hall, N., Jarvis, E. D., Johnson, R. N., Karlsson, E. K., Kress, W. J., Kuraku, S., Lawniczak, M. K. N., Lindblad-Toh, K., Lopez, J. V., … Lewin, H. A. (2022). Why sequence all eukaryotes? Proceedings of the National Academy of Sciences, 119(4), e2115636118. https://doi.org/10.1073/pnas.2115636118

Bovio, E., Garzoli, L., Poli, A., Prigione, V., Firsova, D., McCormack, G. P., & Varese, G. C. (2018). The culturable mycobiota associated with three Atlantic sponges, including two new <i/> species: Thelebolus balaustiformis and T. spongiae. Fungal Systematics and Evolution, 1(1), 141–167. https://doi.org/10.3114/fuse.2018.01.07

Camacho, C., Coulouris, G., Avagyan, V., Ma, N., Papadopoulos, J., Bealer, K., & Madden, T L. (2009). BLAST+: Architecture and applications. BMC Bioinformatics, 10(1), 421. https://doi.org/10.1186/1471-2105-10-421

Cantrell, S. A., Hanlin, R. T., & Emiliano, A. (2007). Periconia variicolor sp. Nov., a new species from Puerto Rico. Mycologia, 99(3), 482–487. https://doi.org/10.1080/15572536.2007.11832573

Chang, Y., Wang, Y., Mondo, S., Ahrendt, S., Andreopoulos, W., Barry, K., Beard, J., Benny, G. L., Blankenship, S., Bonito, G., Cuomo, C., Desiro, A., Gervers, K. A., Hundley, H., Kuo, A., LaButti, K., Lang, B. F., Lipzen, A., O'Donnell, K., … Spatafora, J. W. (2022). Evolution of zygomycete secretomes and the origins of terrestrial fungal ecologies. IScience, 25(8), 104840. https://doi.org/10.1016/j.isci.2022.104840

Crous, P. W., Luangsa-ard, J. J., Wingfield, M. J., Carnegie, A. J., Hernández-Restrepo, M., Lombard, L., Roux, J., Barreto, R. W., Baseia, I. G., Cano-Lira, J. F., Martín, M. P., Morozova, O. V., Stchigel, A. M., Summerell, B. A., Brandrud, T. E., Dima, B., García, D., Giraldo, A., Guarro, J., … Groenewald, J. Z. (2018). Fungal Planet description sheets: 785-867. Persoonia - Molecular Phylogeny and Evolution of Fungi, 41(1), 238–417. https://doi.org/10.3767/persoonia.2018.41.12

Donath, A., Jühling, F., Al-Arab, M., Bernhart, S. H., Reinhardt, F., Stadler, P. F., Middendorf, M., & Bernt, M. (2019). Improved annotation of protein-coding genes boundaries in metazoan mitochondrial genomes. Nucleic Acids Research, 47(20), 10543–10552. https://doi.org/10.1093/nar/gkz833

Espagne, E., Balhadère, P., Penin, M.-L., Barreau, C., & Turcq, B. (2002). HET-E and HET-D Belong to a New Subfamily of WD40 Proteins Involved in Vegetative Incompatibility Specificity in the Fungus Podospora anserina. Genetics, 161(1), 71–81. https://doi.org/10.1093/genetics/161.1.71

Galiana, E., Marais, A., Mura, C., Industri, B., Arbiol, G., & Ponchet, M. (2011). Ecosystem Screening Approach for Pathogen-Associated Microorganisms Affecting Host Disease. Applied and Environmental Microbiology, 77(17), 6069–6075. https://doi.org/10.1128/AEM.05371-11

Galiana, E., Ponchet, M., & Marais, A. (2011). Treatment of plants against oomycete infection (Patent No. WO 2011/110758 A1).

Greiner, S., Lehwark, P., & Bock, R. (2019). OrganellarGenomeDRAW (OGDRAW) version 1.3.1: Expanded toolkit for the graphical visualization of organellar genomes. Nucleic Acids Research, 47(W1), W59–W64. https://doi.org/10.1093/nar/gkz238

Gunasekaran, R., Janakiraman, D., Rajapandian, S. G. K., Appavu, S. P., Namperumalsamy Venkatesh, P, & Prajna, L. (2021). Periconia species—An unusual fungal pathogen causing mycotic keratitis. Indian Journal of Medical Microbiology, 39(1), 36–40. https://doi.org/10.1016/j.ijmmb.2020.10.006

Haas, B. J., Papanicolaou, A., Yassour, M., Grabherr, M., Blood, P. D., Bowden, J., Couger, M. B., Eccles, D., Li, B., Lieber, M., MacManes, M. D., Ott, M., Orvis, J., Pochet, N., Strozzi, F., Weeks, N., Westerman, R., William, T., Dewey, C. N., … Regev, A. (2013). De novo transcript sequence reconstruction from RNA-seq using the Trinity platform for reference generation and analysis. Nature Protocols, 8(8), 1494–1512. https://doi.org/10.1038/nprot.2013.084

Hall, T. A. (1999). BioEdit: A user-friendly biological sequence alignment editor and analysis program for Windows 95/98/NT. Nucleic Acids Symposium Series, 41, 95–98.

Hyde, K., Chaiwan, N., Norphanphoun, C., Boonmee, S., Camporesi, E., Chethana, K., Dayarathne, M., de Silva, N., & Dissanayake, A. (2018). Mycosphere notes 169-224. Mycosphere, 9(2), 271–430. https://doi.org/10.5943/mycosphere/9/2/8

Jones, P., Binns, D., Chang, H.-Y., Fraser, M., Li, W., McAnulla, C., McWilliam, H., Maslen, J., Mitchell, A., Nuka, G., Pesseat, S., Quinn, A. F., Sangrador-Vegas, A., Scheremetjew, M., Yong, S.-Y., Lopez, R., & Hunter, S. (2014). InterProScan 5: Genome-scale protein function classification. Bioinformatics, 30(9), 1236–1240. https://doi.org/10.1093/bioinformatics/btu031

Knapp, D. G., Németh, J. B., Barry, K., Hainaut, M., Henrissat, B., Johnson, J., Kuo, A., Lim, J. H. P., Lipzen, A., Nolan, M., Ohm, R. A., Tamás, L., Grigoriev, I. V., Spatafora, J. W., Nagy, L. G., & Kovács, G. M. (2018). Comparative genomics provides insights into the lifestyle and reveals functional heterogeneity of dark septate endophytic fungi. Scientific Reports, 8(1), 6321. https://doi.org/10.1038/s41598-018-24686-4

Kumar, S., Jones, M., Koutsovoulos, G., Clarke, M., & Blaxter, M. (2013). Blobology: Exploring raw genome data for contaminants, symbionts and parasites using taxon-annotated GC-coverage plots. Frontiers in Genetics, 4. https://doi.org/10.3389/fgene.2013.00237

Laetsch, D. R., & Blaxter, M. L. (2017). BlobTools: Interrogation of genome assemblies. F1000Research, 6, 1287. https://doi.org/10.12688/f1000research.12232.1

Li, H. (2018). Minimap2: Pairwise alignment for nucleotide sequences. Bioinformatics, 34(18), 3094–3100. https://doi.org/10.1093/bioinformatics/bty191

Li, J. Y., Sidhu, R. S., Ford, E. J., Long, D. M., Hess, W. M., & Strobel, G. A. (1998). The induction of taxol production in the endophytic fungus—Periconia sp from Torreya grandifolia. Journal of Industrial Microbiology and Biotechnology, 20(5), 259–264. https://doi.org/10.1038/sj.jim.2900521

Mandyam, K., Fox, C., & Jumpponen, A. (2012). Septate endophyte colonization and host responses of grasses and forbs native to a tallgrass prairie. Mycorrhiza, 22(2), 109–119. https://doi.org/10.1007/s00572-011-0386-y

Manni, M., Berkeley, M. R., Seppey, M., Simão, F. A., & Zdobnov, E. M. (2021). BUSCO Update: Novel and Streamlined Workflows along with Broader and Deeper Phylogenetic Coverage for Scoring of Eukaryotic, Prokaryotic, and Viral Genomes. Molecular Biology and Evolution, 38(10), 4647–4654. https://doi.org/10.1093/molbev/msab199

Marçais, G., & Kingsford, C. (2011). A fast, lock-free approach for efficient parallel counting of occurrences of k-mers. Bioinformatics, 27(6), 764–770. https://doi.org/10.1093/bioinformatics/btr011

Markovskaja, S., & Kačergius, A. (2014). Morphological and molecular characterisation of Periconia pseudobyssoides sp. Nov. And closely related P. byssoides. Mycological Progress, 13(2), 291–302. https://doi.org/10.1007/s11557-013-0914-6

Neuhauser, S., Huber, L., & Kirchmair, M. (2011). A DNA based method to detect the grapevine root-rotting fungus Roesleria subterranea in soil and root samples. 19.

Nicolaisen, M., Frisvad, J. C., & Rossen, L. (1997). A Penicillium freii gene that is highly similar to the β-keto-acyl synthase domain of polyketide synthase genes from other fungi. Letters in Applied Microbiology, 25(3), 197–201. https://doi.org/10.1046/j.1472-765X.1997.00005.x

Nurk, S., Walenz, B. P., Rhie, A., Vollger, M. R., Logsdon, G. A., Grothe, R., Miga, K. H., Eichler, E. E., Phillippy, A. M., & Koren, S. (2020). HiCanu: Accurate assembly of segmental duplications, satellites, and allelic variants from high-fidelity long reads. Genome Research, 30(9), 1291–1305. https://doi.org/10.1101/gr.263566.120

Odvody, G., Dunkle, L., & Edmunds, L. (1977). Phytopathology, 67, 1485-1489. https://doi.org/10.1094/Phyto-67-1485

O’Neil, S. T., Dzurisin, J. D., Carmichael, R. D., Lobo, N. F., Emrich, S. J., & Hellmann, J. J. (2010). RPeoseaprcuh alratictleion-level transcriptome sequencing of nonmodel organisms Erynnis propertius and Papilio zelicaon.

Pellegrin, C., Morin, E., Martin, F. M., & Veneault-Fourrey, C. (2015). Comparative Analysis of Secretomes from Ectomycorrhizal Fungi with an Emphasis on Small-Secreted Proteins. Frontiers in Microbiology, 6. https://doi.org/10.3389/fmicb.2015.01278

Plassart, P., Terrat, S., Thomson, B., Griffiths, R., Dequiedt, S., Lelievre, M., Regnier, T., Nowak, V., Bailey, M., Lemanceau, P., Bispo, A., Chabbi, A., Maron, P.-A., Mougel, C., & Ranjard, L. (2012). Evaluation of the ISO Standard 11063 DNA Extraction Procedure for Assessing Soil Microbial Abundance and Community Structure. PLoS ONE, 7(9), e44279. https://doi.org/10.1371/journal.pone.0044279

Prasher, I. B., & Verma, R. K. (2012). Periconia Species New To North-Western Himalayas. 3.

Ranallo-Benavidez, T. R., Jaron, K. S., & Schatz, M. C. (2020). GenomeScope 2.0 and Smudgeplot for reference-free profiling of polyploid genomes. Nature Communications, 11(1), Article 1. https://doi.org/10.1038/s41467-020-14998-3

Romero, A., Carrion, G., & Rico-Gray, V. (2001). Fungal latent pathogens and endophytes from leaves of. Fungal Diversity, 7.

Sallet, E., Gouzy, J., & Schiex, T. (2019). EuGene: An Automated Integrative Gene Finder for Eukaryotes and Prokaryotes. Methods in Molecular Biology (Clifton, N.J.), 1962, 97–120. https://doi.org/10.1007/978-1-4939-9173-0_6

Schneegurt, M., Dore, S., & Kulpa, C. (2003). Direct Extraction of DNA from Soils for Studies in Microbial Ecology. Current Issues in Molecular Biology. https://doi.org/10.21775/cimb.005.001

Tanaka, K., Hirayama, K., Yonezawa, H., Sato, G., Toriyabe, A., Kudo, H., Hashimoto, A., Matsumura, M., Harada, Y., Kurihara, Y., Shirouzu, T., & Hosoya, T. (2015). Revision of the Massarineae (Pleosporales, Dothideomycetes). Studies in Mycology, 82(1), 75–136. https://doi.org/10.1016/j.simyco.2015.10.002

Taylor, T. N., Krings, M., & Taylor, E. L. (2015). Ascomycota. In Fossil Fungi (pp. 129–171). Elsevier. https://doi.org/10.1016/B978-0-12-387731-4.00008-6

Teufel, F., Almagro Armenteros, J. J., Johansen, A. R., Gíslason, M. H., Pihl, S. I., Tsirigos, K. D., Winther, O., Brunak, S., von Heijne, G., & Nielsen, H. (2022). SignalP 6.0 predicts all five types of signal peptides using protein language models. Nature Biotechnology, 40(7), 1023–1025. https://doi.org/10.1038/s41587-021-01156-3

Tischer, M., Gorczak, M., Bojarski, B., Paw?owska, J., Hoffeins, C., Hoffeins, H. W., & Wrzosek, M. (2019). New fossils of ascomycetous anamorphic fungi from Baltic amber. Fungal Biology, 804–810.

Verma, V. C., Lobkovsky, E., Gange, A. C., Singh, S. K., & Prakash, S. (2011). Piperine production by endophytic fungus Periconia sp. Isolated from Piper longum L. The Journal of Antibiotics, 64(6), 427–431. https://doi.org/10.1038/ja.2011.27

Vilgalys, R., & Hester, M. (1990). Rapid genetic identification and mapping of enzymatically amplified ribosomal DNA from several Cryptococcus species. Journal of Bacteriology, 172(8), 4238–4246. https://doi.org/10.1128/jb.172.8.4238-4246.1990

White, T. J., Bruns, T., Lee, S., & Taylor, J. (1990). AMPLIFICATION AND DIRECT SEQUENCING OF FUNGAL RIBOSOMAL RNA GENES FOR PHYLOGENETICS. In PCR Protocols (pp. 315–322). Elsevier. https://doi.org/10.1016/B978-0-12-372180-8.50042-1

Yang, E.-F., Phookamsak, R., Jiang, H.-B., Tibpromma, S., Bhat, D. J., Karunarathna, S. C., Dai, D.-Q., Xu, J.-C., & Promputtha, I. (2022). Taxonomic Reappraisal of Periconiaceae with the Description of Three New Periconia Species from China. Journal of Fungi, 8(3), 243. https://doi.org/10.3390/jof8030243

Zheng, F., Han, T., Basit, A., Liu, J., Miao, T., & Jiang, W. (2022). Whole-Genome Sequence and Comparative Analysis of Trichoderma asperellum ND-1 Reveal Its Unique Enzymatic System for Efficient Biomass Degradation. Catalysts, 12(4), 437. https://doi.org/10.3390/catal12040437

Zhu, Y., Dong, J., Wang, L., Zhou, W., Li, L., He, H., Liu, H., & Zhang, K. (2008). Screening and isolation of antinematodal metabolites againstBursaphelenchus xylophilus produced by fungi. Annals of Microbiology, 58(3), 375–380. https://doi.org/10.1007/BF03175531

Sallet, E., Gouzy, J., & Schiex, T. (2019). EuGene : An Automated Integrative Gene Finder for Eukaryotes and Prokaryotes. In M. Kollmar (Éd.), Gene Prediction (Vol. 1962, p. 97–120). Springer New York. doi: 10.1007/978-1-4939-9173-0_6

UniProt Consortium, T. (2018). UniProt: The universal protein knowledgebase. Nucleic Acids Research, 46(5), 2699–2699. doi: 10.1093/nar/gky092

